# Distinct cellular and transcriptional mechanisms mediate an antioxidant therapeutic response in 22q11-deleted upper layer cortical projection neurons

**DOI:** 10.64898/2025.12.03.692138

**Authors:** Shah Rukh, Daniel W. Meechan, Abra Roberts, Connor Siggins, Zachary D. Erwin, Thomas M. Maynard, Anthony-S. LaMantia

## Abstract

We characterized cellular and molecular mechanisms underlying the therapeutic response *in vitro* and *in vivo* to the antioxidant N-acetyl cysteine (NAC), which in the *LgDel* 22q11.2 Deletion Syndrome mouse model restores growth and connectivity of upper layer cortical projection neurons (<u>L</u>ayer 2/3 PNs) and improves cognitive performance. NAC treatment of primary cultured *LgDel* L 2/3 PNs does not restore these neurons to a wild type (WT) state. Rather than returning to the bimodal dendrite and axon size distribution seen in WT, *LgDel* L 2/3 PN dendrite and axon growth *in vitro* increases unimodally in response to NAC. In parallel, altered expression of 22q11-deleted genes and presumed downstream targets are unchanged. Instead, novel antioxidant defense and neuronal growth genes are differentially expressed: some generally NAC-regulated, others responsive only in the context of 22q11 deletion. Apparently, NAC ameliorates L 2/3 PN developmental pathology without restoring WT cell states or typical expression of mutant genes or their downstream targets. NAC also elicits differential expression of antioxidant defense genes in 22q11-deleted L 2/3 PNs—but not L 5/6 counterparts—in the developing postnatal *LgDel* mouse cortex, rather than modulating 22q11 genes or downstream targets. These NAC-dependent, L 2/3 PN-selective *in vivo* cellular and transcriptional changes differ substantially from those in primary culture. Thus, despite some *in vitro* and *in vivo* parallels, the NAC therapeutic response that diminishes oxidative stress-related L 2/3 PN circuit and behavioral pathology due to 22q11 deletion has a unique *in vivo* signature.

## INTRODUCTION

Therapeutic approaches for neurodevelopmental disorders (NDDs) with known genetic causes have focused on restoring function of single mutant genes in monogenic disorders or one or more critical mutant genes in polygenic disorders (Berry-Kravis et al., 2018; Chen and Geschwind, 2022; Devinsky et al., 2025; Lauffer et al., 2024; Sandweiss et al., 2020; Zylka, 2020). Nevertheless, substantial challenges in targeting, timing, and regulation of functional copies of mutant genes or RNA therapies remain (Bajikar et al., 2024). Alternately, downstream genes or networks disrupted by NDD-associated mutations are considered prime therapeutic targets. Nevertheless, aligning these targets with effective therapeutic agents remains difficult (Copping et al., 2021; Diaz-Caneja et al., 2021; Paul and Potter, 2024; Pedini et al., 2023). To determine whether restoring mutant or downstream target gene function is necessary for therapeutic efficacy, we asked whether N-acteyl cysteine (NAC), an antioxidant that ameliorates <u>L</u>ayer 2/3 Projection Neuron (L 2/3 PNs) pathology—a key pathological feature of multiple NDDs (Batiuk et al., 2022; Lewis and Sweet, 2009; Parikshak et al., 2013; Sacai et al., 2020)—and related behavioral deficits in the *LgDel* mouse model of 22q11.2 Deletion Syndrome (22q11; Meechan et al., 2015a) restores L 2/3 PN cellular, metabolic and transcriptional states directly altered by 22q11 deletion or elicits a divergent therapeutic response, independent of 22q11 gene dosage-mediated changes in L 2/3 PNs *in vitro* as well as *in vivo*.

22q11DS, one of the most common polygenic/copy number variant developmental syndromes, also carries one of the most robust genetic risks for Schizophrenia, Autistic Spectrum Disorder and other psychiatric/NDD pathologies. Thus, therapeutic approaches that target specific neurons and circuits compromised in 22q11DS may provide a template for parallel interventions in a broader range of NDDs. It is not clear, however, how to integrate mechanisms inferred from *in vitro* assays with *in vivo* actions of any potential therapeutic agent in 22q11DS or any other NDD. Human 22q11-deleted induced pluripotent stem cells (IPSCs) have yet to generate neurons or glia matched to *in vivo* classes compromised by 22q11 deletion pathology. Thus, this promising approach has not yet securely identified discrete targets for therapeutic development (Fiksinski et al., 2023; LaMarca et al., 2018; Lin et al., 2016; Rao et al., 2025a). We showed previously in the *LgDel* mouse that NAC selectively reverses oxidative stress in L 2/3 PNs *in vitro* and *in vivo*, apparently by diminishing elevated reactive oxygen species (ROS) and reversing their impact on L2/3 PN growth as well as cytological and synaptic integrity (Fernandez et al., 2019). It is not clear, however, whether NAC returns 22q11-deleted L 2/3 PNs *in vitro* or *in vivo* to typical cell biological, molecular, or transcriptional states. Indeed, NAC may establish a divergent “therapeutic” state via cell biological, metabolic and transcriptional responses that circumvent changes caused by diminished 22q11 gene dosage.

We resolved these issues using *in vitro* and *in vivo* assays to define growth, metabolic, and transcriptional states in WT and *LgDel* L 2/3 PNs with and without NAC treatment, as well as those in a related heterozygous mutation of key 22q11-deleted mitochondrial gene, *Txnrd2*, also associated with ROS-mediated, NAC-sensitive L 2/3 PN pathology. NAC ameliorates, but does not directly reverse, cell growth, metabolic and transcriptional disruptions due to diminished dosage of multiple 22q11 genes in differentiating L 2/3 PNs. Apparently, circumventing changes due to aberrant cellular states like oxidative stress rather than directly restoring mutant or downstream gene function can provide significant—and perhaps more readily achievable— therapeutic benefits in genetically and phenotypically complex NDDs like 22q11DS.

## RESULTS

### A distinct transcriptional state defines 22q11 gene deleted L 2/3 PNs in vitro

We first analyzed L 2/3 PNs *in vitro* (*iv*) from mouse E16.5 *LgDel* cortices since they provide a uniform population for cellular and transcriptome comparison, and their pathological changes parallel those of *LgDel* L 2/3 PNs *in vivo* (Fernandez et al, 2019). We confirmed limited growth for *LgDel* L 2/3 PNs *iv* (**Figure 1**), and assessed the distribution of L 2/3 PN *iv* sizes, based upon total dendrite or axon length per cell. WT L 2/3 PN *iv* dendritic arbors are bimodally distributed (**Figure 1B**, *top*; see also **Figure 3**). In contrast, *LgDel* L 2/3 PNs *iv* dendritic arbors are distributed around a single mean that overlaps the smaller WT population. In parallel, the *LgDel* L 2/3 PN *iv* axon distribution (**Figure 1B**, *bottom*) lacks the WT “tail” of larger axons. Finally, the modest but significant correlation between dendrite and axon length for individual WT L 2/3 PNs *iv* (r^2^ = 0.1345, p = 0.004; **Supp. Fig. 1**) is lost in *LgDel* (r^2^ = 0.0065, p = 0.616). Thus, 22q11 gene deletion either eliminates a population of L 2/3 PN *iv* neuroblasts with maximal dendrite and axon growth capacity or limits growth capacities of a subset of L 2/3 PNs *iv*.

**Figure 1:**
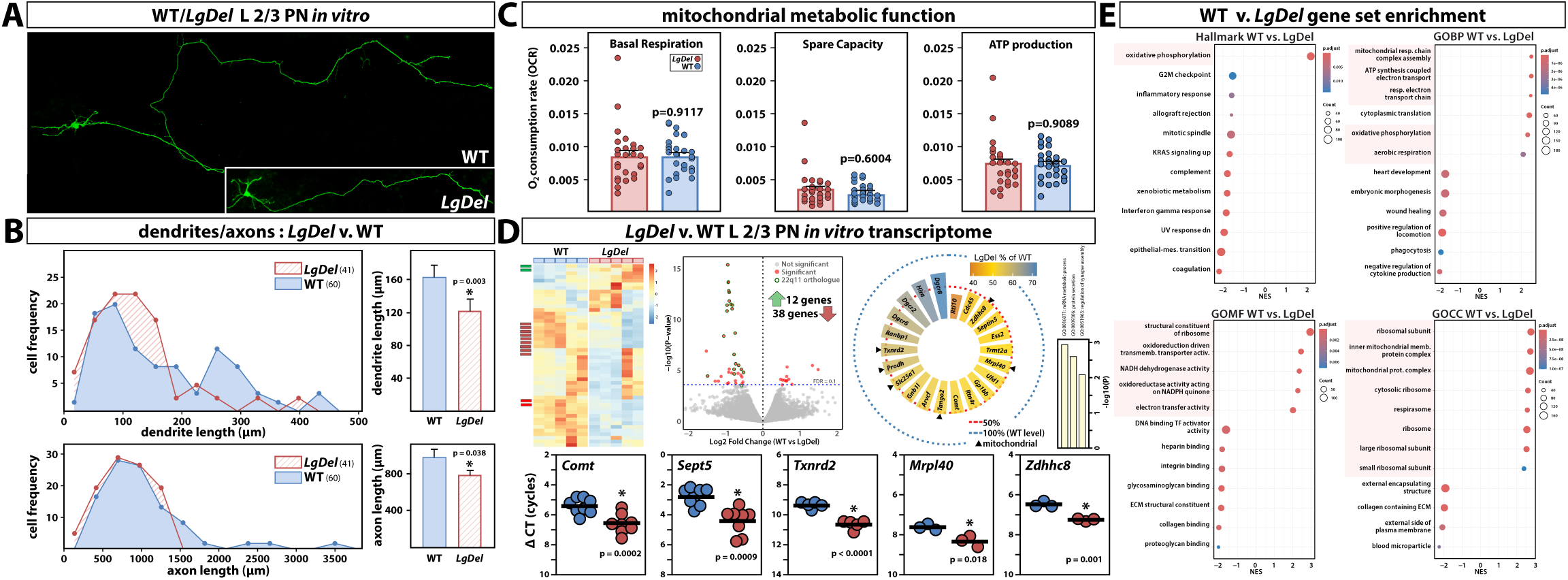
Distinct metabolic and transcriptional states accompany selective dendrite and axon growth changes due to 22q11 gene deletion in mouse primary L 2/3 PNs iv. **A**. Distinctions between WT (*top*) and *LgDel* (*bottom*, *<u>inset</u>*) L 2/3 PNs *iv* with larger vs. smaller dendrite and axon arbors. **B**. Frequency distribution of WT and *LgDel* L 2/3 PNs *iv* based upon total dendrite (*top*, *left*) or axon (*bottom*, *left*) length (n = 55 WT, 36 *LgDel* L 2/3 PNs; one way ANOVA) and mean differences in total dendrite (*top*, *right*) and axon lengths (*bottom*, *right*). **C**. Basal respiration, spare capacity and ATP production in *LgDel* vs. WT littermate L 2/3 PNs *iv* based upon O_2_ consumption (Seahorse Metabolic Analyzer; n= 19 *LgDel*, 16 WT, multi-well cultures across 4 litters; one way ANOVA). **D**. Comparison of WT and *LgDel* L 2/3 PNs *iv* transcriptomes based upon 5 biological replicates of pooled L 2/3 PN samples (n= 15 fetuses/6 litters WT; 15 fetuses/7 litter *LgDel*) identifies 50 differentially expressed (DE) genes in *LgDel* L 2/3 PNs (FDR < 0.1), including 21 heterozygous-deleted 22q11 orthologues (*top*, *right*). A gene ontology (GO) analysis of 19 non-22q11 deleted genes (*top*, *far right*) indicates DE genes are enriched for metabolic and neuron-specific pathways. qPCR analysis of parallel RNA samples from WT and *LgDel* L 2/3 PNs *iv* confirms 50% significantly diminished expression of 5 22q11deleted genes. **E**. Hallmark gene set enrichment analysis (GSEA) and gene ontologies (GO) associated with biological processes (GOBP), molecular functions (GOMF) and cellular compartments (GOCC) identify enrichment of gene sets associated with oxidative phosphorylation and mitochondrial function in WT vs. *LgDel* L 2/3 PNs.

**Figure 2:**
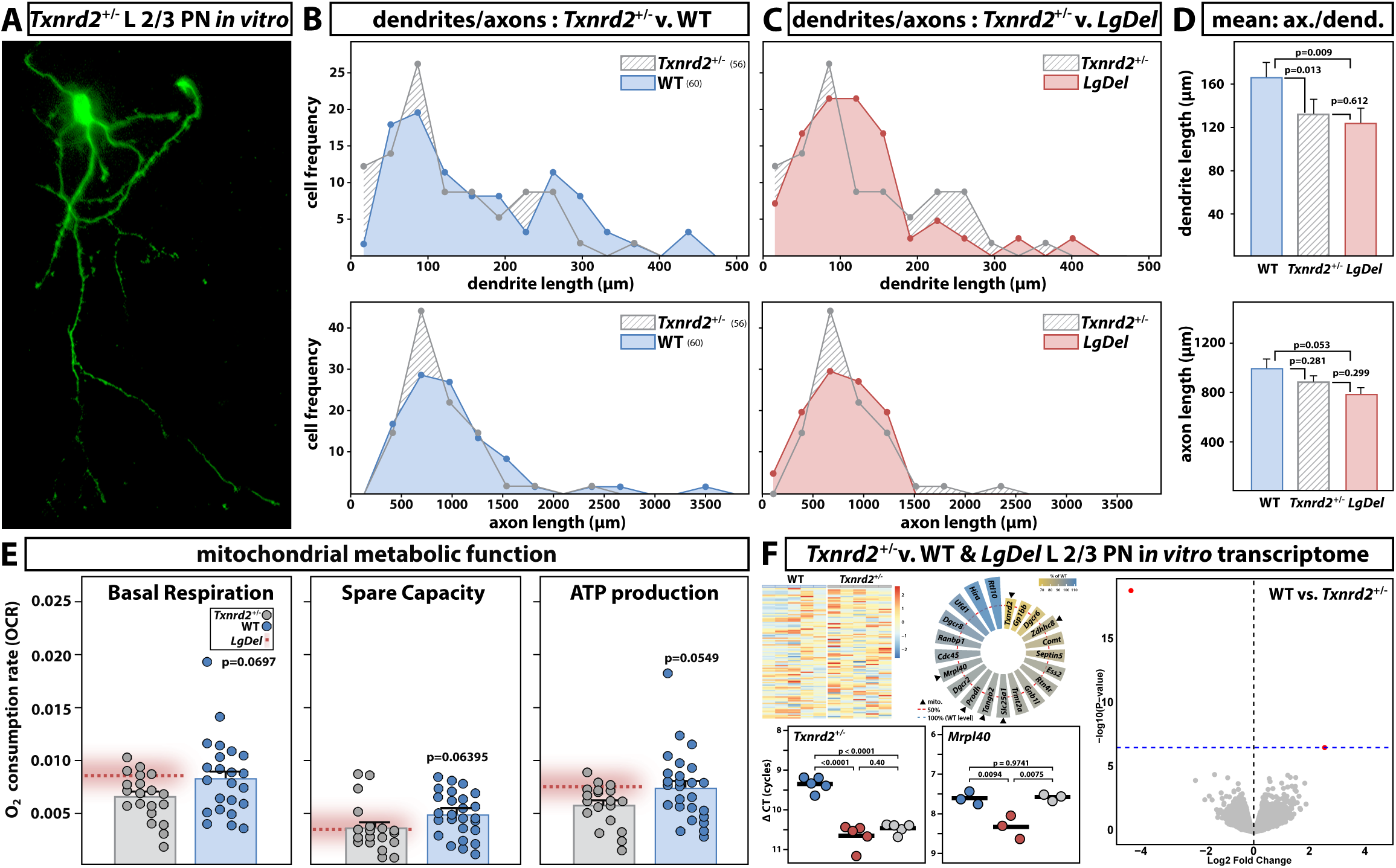
Distinct growth, metabolic and transcriptional responses of Txnrd2^+/-^ L 2/3 PNs iv. **A**. An example of a L 2/3 PN *iv* grown from an E16.5 electroporated cortical neuroblast harvested from a constitutive heterozygous deleted *Txnrd2* mutant fetus. **B**. Bimodal frequency distributions of smaller vs. larger dendrite arbors based upon total dendrite length in WT and Txnrd2+/- L 2/3 PNs iv. The *Txnrd2*^+/-^ distribution, however, is shifted slightly leftward. **C**. Similar frequency distributions of Txnrd2+/- and WT axon lengths accompanied increased frequency of smaller axons, and diminished frequency of larger axons in *Txnrd2*^+/-^ L 2/3 PNs iv. **D**. Comparison of the mean dendrite and axon lengths in WT, *Txnrd2*^+/-^ and LgDel L 2/3 PNs iv (n= 55 WT, 51 Txnrd2+/-, 36 LgDel L 2/3 PNs iv; one way ANOVA). **E**. Mitochondrial basal respiration, spare capacity and ATP production differ in Txnrd2+/- vs. WT littermate control cultures (calculated from O_2_ consumption; Agilent Seahorse; n= 26 WT, 19 *Txnrd2*^+/-^ cultures across 6 litters; T-test). **F**. Few significant differences, including significantly DE genes, in *Txnrd2*^+/-^ vs WT L 2/3 PN transcriptomes (*top*; n= 15 fetuses/5 litter *Txnrd2*^+/-^; 15 fetuses/6 litters WT). Two novel DE genes are distinct from the DE gene set identified in the *LgDel* vs. WT comparison. In addition, *Txnrd2* expression is diminished in *Txnrd2*^+/-^ L 2/3 PNs *iv*, validated by qPCR (*bottom*).

**Figure 3:**
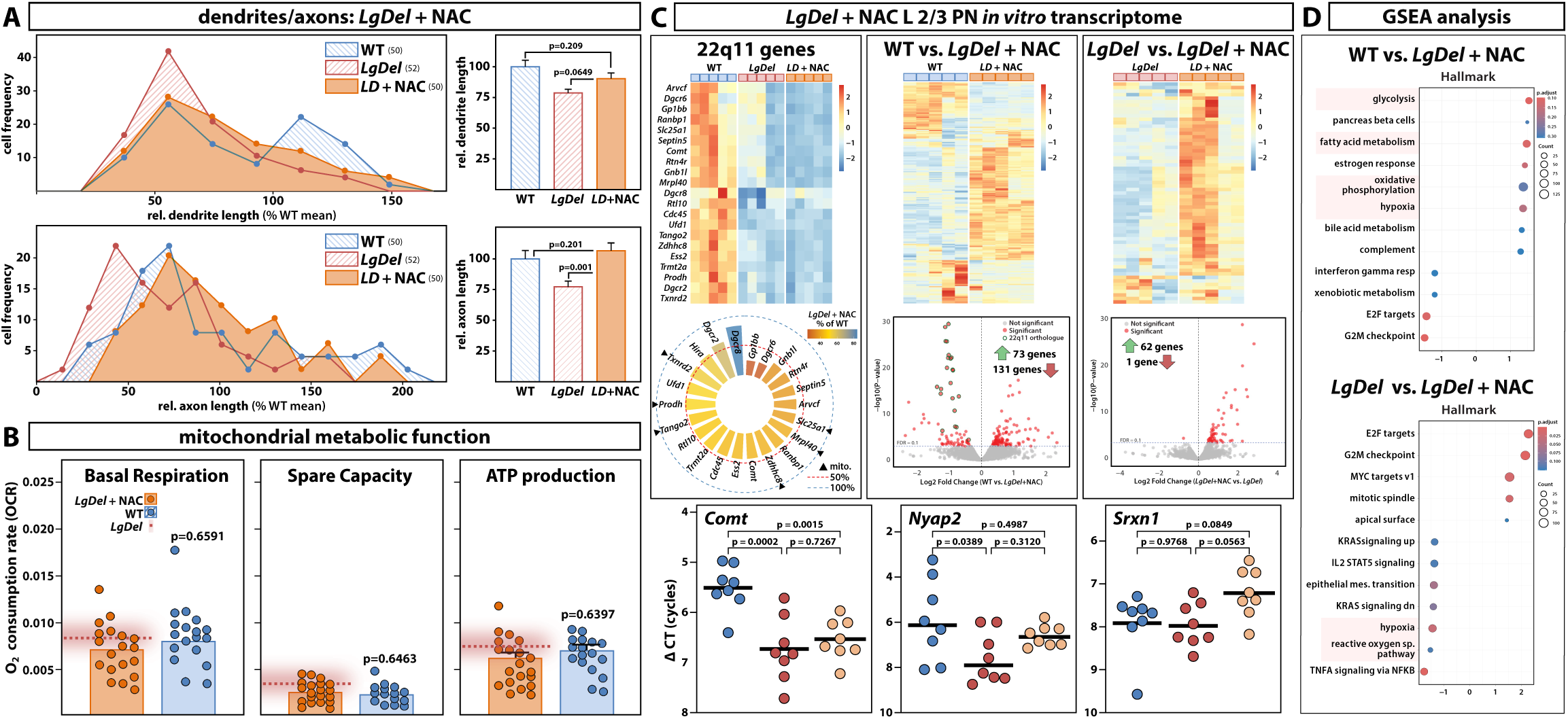
Distinct cellular and transcriptional therapeutic responses to N-acetyl cysteine (NAC) increase L 2/3 PN iv dendrite and axon growth. **A**. NAC treatment yields a distinct unimodally distributed population of L 2/3 PNs iv with a mean increase in dendrite lengths that does not reflect return to the WT bimodal distribution (*top*, *left*; *data reanalyzed from* Fernandez et al, 2019). Mean differences in dendrite length in *LgDel* + NAC L 2/3 PNs iv are statistically indistinguishable from WT and *LgDel* (*top*, *right*). NAC restores the distribution of axon lengths toward the WT distribution (*bottom*, *left*), and fully returns mean lengths to a value statistically indistinguishable from WT (*bottom*, *right*). **B**. *LgDel* + NAC L 2/3 PNs have basal respiration, spare capacity and ATP production levels statistically indistinguishable from WT littermate controls (n= 17 cultures from 4 litters, one way ANOVA), and comparable to *LgDel* L 2/3 PNs *iv* (see **Figure 1C**). **C**. Transcriptome analysis of *LgDel* + NAC L 2/3 PNs *iv*. The 21 22q11 genes for which transcripts are detected above threshold in WT and *LgDel* L 2/3 PNs are diminished by 50% in *LgDel* + NAC L 2/3 PNs (*top*, *left*) as in *LgDel* counterparts. The transcriptomes of *LgDel* + NAC L 2/3 PNs differ from WT counterparts: there are 204 DE genes (*top*, *middle*) (n= 15 fetuses/6 litters WT; 15 fetuses/7 litter *LgDel* + NAC). Of 131 down-regulated DE genes, 21 are 22q11 orthologues deleted in *LgDel*. When *LgDel* + NAC L 2/3 PN *iv* transcriptome is compared to *LgDel*, none of the 64 DE genes overlap DE genes identified in the WT vs. *LgDel* comparison (n= 15 fetuses/7 litters *LgDel*; 15 fetuses/7 litter LgDel + NAC). Validation of a novel antioxidant defense gene (*bottom*, *left*), *Srxn1*, based upon qPCR assessement. qPCR expression analysis of a representative *LgDel* vs. WT DE gene, *Nyap2*, which is significantly down-regulated when *LgDel* L 2/3 PNs *iv* are compared to WT, but does not change significantly from WT levels in *LgDel* + NAC L 2/3 PNs *iv* (*bottom*, *middle*). **D**. Hallmark GSEA analysis of WT vs. *LgDel* + NAC (*top*) and *LgDel* vs. *LgDel* + NAC (*bottom*) whole transcriptomes. The WT vs. *LgDel* + NAC comparison identifies positive enrichment multiple gene sets associated with mitochondrial function (orange highlight; *top*). The *LgDel* vs. *LgDel* + NAC comparison identifies negative enrichment for hypoxia and ROS pathway gene sets (*red shading*), consistent with NAC antioxidant function.

We next asked whether growth-restricted *LgDel* vs. WT L 2/3 PNs *in vitro* have altered mitochondrial function assessed via O_2_ consumption. We found no significant mean differences in 3 major domains of mitochondrial metabolism: basal respiration, spare capacity, and ATP production (**Figure 1C**). Apparently, *LgDel* L 2/3 PNs *in vitro* retain aspects of typical mitochondrial and metabolic function, despite selectively limited dendrite and axon growth as well as elevated ROS (Fernandez et al., 2019). Stable metabolism in 22q11-deleted L 2/3 PNs may reflect a mitochondrial homeostatic response in the context of an altered redox environment (Panieri and Santoro, 2016; Shadel and Horvath, 2015; Zhang et al., 2022).

A distinct transcriptional state accompanies selectively restricted growth and apparent mitochondrial homeostasis in *LgDel* L 2/3 PNs *iv* (**Figure 1D**, *top row, left*). 50 genes are significantly differentially expressed (DE; FDR < 0.1): 38 diminished and 12 enhanced (**Figure 1D**, *top row, center*). Twenty-one 22q11 deleted genes, including all 6 mitochondria-localized transcripts (Campbell et al., 2023; Maynard et al., 2008; Motahari et al., 2019) are among the 38 down-regulated DE genes. qPCR of a subset of 22q11 genes confirms the 50% decrease in L 2/3 PNs *iv* (**Figure 1D**, *bottom row*). The 17 remaining down-regulated DE genes include: *Nyap2* (Yokoyama et al., 2011) and *Gucy1a2* (Haase et al., 2010), associated with neuron growth; *Pclo* and *Thbs2* (Mazur et al., 2021; Waites et al., 2013) implicated in synapse formation and maintenance; *Proser1* (Fleming et al., 2024) a chromatin regulator; *Lnpep* (Zhang et al., 2017), a contributor to synaptic plasticity, and *Vps13c* (Monfrini et al., 2022; Schrӧder et al., 2024) a modulator of mitochondrial homeostasis. Significantly up-regulated transcripts include: *Emc10*, a regulator of neural development (Shao et al., 2021) previously associated with 22q11 deletion (Thakur et al., 2025); *Aopep* which modulates synaptogenesis (Garavaglia et al., 2022), and *B9d2*, associated with cilia signaling (Caenen-Braz et al., 2024). A parallel gene ontology (GO) analysis indicates that DE genes engage mRNA metabolism, protein secretion and synapse assembly gene sets (**Figure 1D**, *far right*). qPCR confirms differential expression of *Nyap2* (see **Figure 3**); however, 2 other DE genes, *Lnpep* and *Gucy1a2* do not differ significantly (*not shown*). Thus, DE genes that modulate bioenergetic homeostasis, neuron or synaptic differentiation in L 2/3 PNs *iv* may contribute to altered growth and differentiation of 22q11-deleted L 2/3 PNs *iv*.

To further analyze divergence between the full transcriptomes of WT (20,650 transcripts) and *LgDel* (25,200 transcripts) L 2/3 PNs *iv*, we used Gene Set Enrichment Analysis (GSEA, **Figure 1E***, top left*) (Subramanian et al., 2005). The most significantly positively enriched Hallmark gene set in *LgDel* L 2/3 PNs *iv* is oxidative phosphorylation, consistent with transcriptional modulation of mitochondrial homeostasis due to oxidative stress. Additional GSEA approaches identify significant positive enrichment in metabolism-related gene sets (**Figure 1E**, *top right* & *bottom*). This positive enrichment of metabolic/homeostasis gene sets was matched by depleted gene sets associated with cytoskeletal dynamism, cell adhesion, and motility. Apparently, 22q11deleted L 2/3 PNs *iv* acquire a transcriptional state that supports metabolic and mitochondrial homeostasis while limiting neuronal growth and differentiation.

### Constitutive Txnrd2 heterozygous deletion and L 2/3 PN iv growth regulation

Acutely diminished expression of the 22q11 mitochondrial gene *Txnrd2*, a rate limiting enzyme for mitochondrial ROS clearance (Prasad et al., 2014; Soerensen et al., 2008) *in vitro* or *in vivo* limits L 2/3 PNs dendrite and axon growth similar to broader 22q11 gene deletion. To determine whether <u>constitutive</u> *Txnrd2* heterozygous deletion accounts for limited L 2/3 PN *iv* dendrite and axon growth in the context of broader 22q11 deletion, we cultured L 2/3 PNs from E16.5 *Txnrd2*^+/-^ fetuses (**Figure 2**). *Txnrd2*^+/-^ L 2/3 PN *iv* dendrite lengths are bimodally distributed (**Figure 2B**, *top*); however, there is a uniform leftward shift (**Figure 2C**, *top*), resulting in an intermediate mean dendrite length between WT and *LgDel*—significantly different from WT, but not *LgDel* (**Figure 2D**, *top*). *Txnrd2*^+/-^ L 2/3 PNs iv axon lengths are distributed similarly to WT (**Figure 2B**, *bottom*) rather than *LgDel* (**Figure 2C**, *bottom*); however, there is less of a “tail” of larger axons. *Txnrd2*^+/-^ axon lengths also shift leftward (**Figure 2C**, *left*), resulting in a non-significant decline in mean length between WT and *LgDel* values (**Figure 2D**, *bottom*). Thus, unlike acutely diminished *Txnrd2* expression *in vitro* and *in vivo* (Fernandez et al., 2019), constitutive heterozygous *Txnrd2* deletion does not substantially modify the distribution of L 2/3 PN *iv* dendrite or axon lengths, or significantly reduce L 2/3 PN *iv* axon growth.

Based upon apparent *Txnrd2* function in mitochondrial ROS clearance in L 2/3 PNs (Fernandez et al., 2019; Maynard et al., 2008), constitutive *Txnrd2* heterozygous deletion might alter L 2/3 PN *iv* mitochondrial metabolism as part of a compensatory mechanism. Indeed, L2/3 PN *iv* O_2_ consumption measurements diverge in *Txnrd2*^+/-^ vs. WT littermate control L 2/3 PNs *iv* (**Figure 2E**), unlike parallel measurements in *LgDel* and WT. Basal respiration, spare capacity and ATP production are all lower in *Txnrd2*^+/-^ L 2/3 PNs *iv*. Differences for basal respiration and spare capacity approach statistical significance, and that for ATP production is marginally significant (p = 0.0697; 0.0639, and 0.0549, respectively). Accordingly, mitochondrial function in the context of constitutive *Txnrd2* heterozygous deletion diverges from that in WT as well as L 2/3 PNs *iv* with full deletion of 22q11 orthologues including *Txnrd2*.

We next asked whether transcriptome changes similar to those in *LgDel* L 2/3 PNs *iv* accompany differences in *Txnrd2*^+/-^ L 2/3 PN *iv* growth and mitochondrial function. With the exception of significantly diminished expression of *Txnrd2* itself, the *Txnrd2*^+/-^ transcriptome does not change in parallel with that of *LgDel* L 2/3 PNs *iv* (**Figure 2F**); instead, it is similar to WT. *Txnrd2* expression, assessed by qPCR, declines by approx. 50% in L 2/3 PNs *iv*, similar to *LgDel*; however, expression levels of other 22q11 deleted genes, including the other 5 mitochondrial genes, do not change (**Figure 2F**, *left*). Finally, only two transcripts met criterion as DE genes compared to WT. *GM8515*, which encodes a predicted protein of unknown function, expressed at low levels in *Txnrd2*^+/-^ and WT L 2/3 PNs, is down-regulated, and *Saa2*, a serum amyloid protein expressed at variable levels, is modestly up-regulated (**Figure 2F**, *right*). Neither, however, is differentially expressed in *LgDel* L2/3 PNs *iv*. Thus, constitutive *Txnrd2* heterozygous deletion does not substantially alter the transcriptional state of L 2/3 PNs *iv*.

Finally, to fully assess transcriptional changes independent of DE genes, we compared the full transcriptomes of *Txnrd2*^+/-^ L 2/3 PNs *iv* (21,774 transcripts) to WT counterparts using Gene Set Enrichment Analysis. The Hallmark comparison between WT and *Txnrd2*^+/-^ transcriptomes yielded no significantly enriched gene sets, while that between *LgDel* and *Txnrd2*^+/-^ yielded enrichment in *LgDel* that parallels *LgDel* vs. WT (see **Figure 1E**). Thus, constitutive heterozygous deletion and reduced expression of *Txnrd2* limits growth and modifies metabolic state of L 2/3 PNs *iv* without substantial transcriptional changes, unlike broader 22q11 deletion.

### 22q11-deleted L 2/3 PN iv growth & transcriptional response to NAC treatment

The ROS scavenger n-Acetyl Cysteine (NAC), is one of a small group of mitochondria-targeted compounds that modifies oxidative stress-related changes in 22q11DS animal models, human 22q11 gene-deleted IPSCs, and individuals with 22q11DS (Crockett et al., 2025; Li et al., 2019). In *LgDel* L 2/3 PNs *iv* NAC diminishes mitochondrial and cytosolic ROS, restores axon/dendrite growth and branching; in *LgDel* L 2/3 PN *in vivo,* it enhances dendrite growth, reduces cytological signs of oxidative stress, increases synapse frequency and restores typical synaptic morphology (Fernandez et al., 2019). We re-analyzed previous neuron growth data from WT & *LgDel* L 2/3 PNs *iv* cultured with and without NAC and confirmed the bimodal distribution of WT L 2/3 PN *iv* dendrite and axon lengths (see **Figure 1**), as well as the unimodal distribution for *LgDel* L 2/3 PNs *iv* (**Figure 3A**). NAC does not fully restore *LgDel* L 2/3 PN dendrite or axon lengths to their WT distribution. Instead, dendrite lengths are monotonically distributed across larger values (**Figure 3A**, *top*) and axon lengths approximate WT (**Figure 3A**, *bottom*). Finally, a modest but significant dendrite/axon size correlation, less robust than for that in WT, returns for *LgDel* + NAC L 2/3 PNs *iv* (**Supp. Fig. 1**). In parallel, mitochondrial metabolic function does not change from values in WT or *LgDel* L 2/3 PNs (**Figure 3B**). This suggests that NAC maintains mitochondrial homeostasis as a part of a therapeutic adjustment in that returns 22q11-deleted L 2/3 PNs *iv* to a state of enhanced—but not fully typical—growth.

NAC-dependent restoration of *LgDel* L 2/3 PN *iv* growth may return *LgDel* DE genes to WT levels, or elicit a unique transcriptional response independent of 22q11 genes or their downstream targets. Despite increased dendrite/axon growth in response to NAC, all 21 of the 22q11-deleted genes expressed in *LgDel* L 2/3 PNs *iv* remain significantly diminished (**Figure 3C**, *top, left & bottom, left*). The transcriptional state of *LgDel* L 2/3 PNs + NAC resembles neither that in WT nor *LgDel* (**Figure 3C**). *LgDel* + NAC L 2/3 PNs *iv* acquire a distinct transcriptional state. 63 genes that are differentially expressed between *LgDel* vs. *LgDel* + NAC L 2/3 PNs *iv* do not include any of the 29 non-22q11 deleted genes that distinguish *LgDel* L 2/3 PNs *iv* from WT (**Figure 3C**, *right*). Instead, NAC-dependent DE genes include 2 up-regulated transcripts associated with Nrf2-regulated antioxidant defense: (**Figure 3C**, *right*; Brennan et al., 2017; Zhou et al., 2015). Thus, a distinct transcriptional response, rather than restoration of WT expression levels of genes altered by 22q11 gene deletion, underlies enhanced dendrite and axon growth in 22q11 gene-deleted L 2/3 PNs *iv* following NAC antioxidant treatment.

Hallmark GSEA analyses of the broader *LgDel* + NAC L 2/3 PN *iv* (19730 transcripts) transcriptomes reinforce the impression that NAC elicits a transcriptional state that supports metabolic homeostasis and mitochondrial function in 22q11-deleted L 2/3 PNs *iv* (**Figure 3D**, *top*). Only one gene set, for oxidative phosphorylation, overlaps WT vs. *LgDel* (see **Figure 1E**), albeit with lower significance. The *LgDel* vs. *LgDel* + NAC comparison identifies negative enrichment for hypoxia and ROS pathway gene sets (**Figure 3D**, *bottom*), consistent with NAC antioxidant function relieving transcriptional correlates of oxidative stress in *LgDel* L 2/3 PNs *iv*. In addition, positive enrichment of E2F targets and G2M checkpoint gene sets (negatively enriched in the *LgDel* vs. WT comparison, see **Figure 1E**) is consistent with NAC acting reversing cellular stress that restricts growth or biases cells toward apoptosis. Apparently, NAC supports “therapeutic” L 2/3 PNs *iv* growth in the context of a novel transcriptional state that modulates antioxidant defense and diminishes effects of 22q11 deletion-dependent cellular stress.

### L 2/3 PNs have 22q11 deletion-specific growth and transcriptional responses to NAC

The unique transcriptional response in NAC-treated 22q11 gene-deleted *LgDel* L 2/3 PNs *iv* may encompass all or a subset of genes also regulated by NAC in WT L 2/3 PNs *iv*. Our previous observations, however, indicate that NAC limits rather than enhances, WT L 2/3 PN *iv* dendrite and axon growth (Fernandez et al., 2019). Indeed, the distribution of dendrite lengths in WT L 2/3 PNs + NAC *iv* resembles that in *LgDel* (**Figure 4A**, *top left*). Mean length is modestly, but significantly lower than *LgDel* (**Figure 4A**, *top right*). NAC has a distinct impact on WT axon growth: axon sizes are distributed around a single mean, lower than the 3 other conditions (**Figure 4A**, *bottom left*). Nevertheless, NAC does not appear to alter mitochondrial metabolism in WT L 2/3 PNs; basal respiration, spare capacity or ATP production are statistically indistinguishable from WT, *LgDel* and *LgDel* + NAC (**Figure 4B**). Apparently, NAC limits WT L 2/3 PN *iv* growth and differentiation without disrupting key dimensions of mitochondrial function.

**Figure 4:**
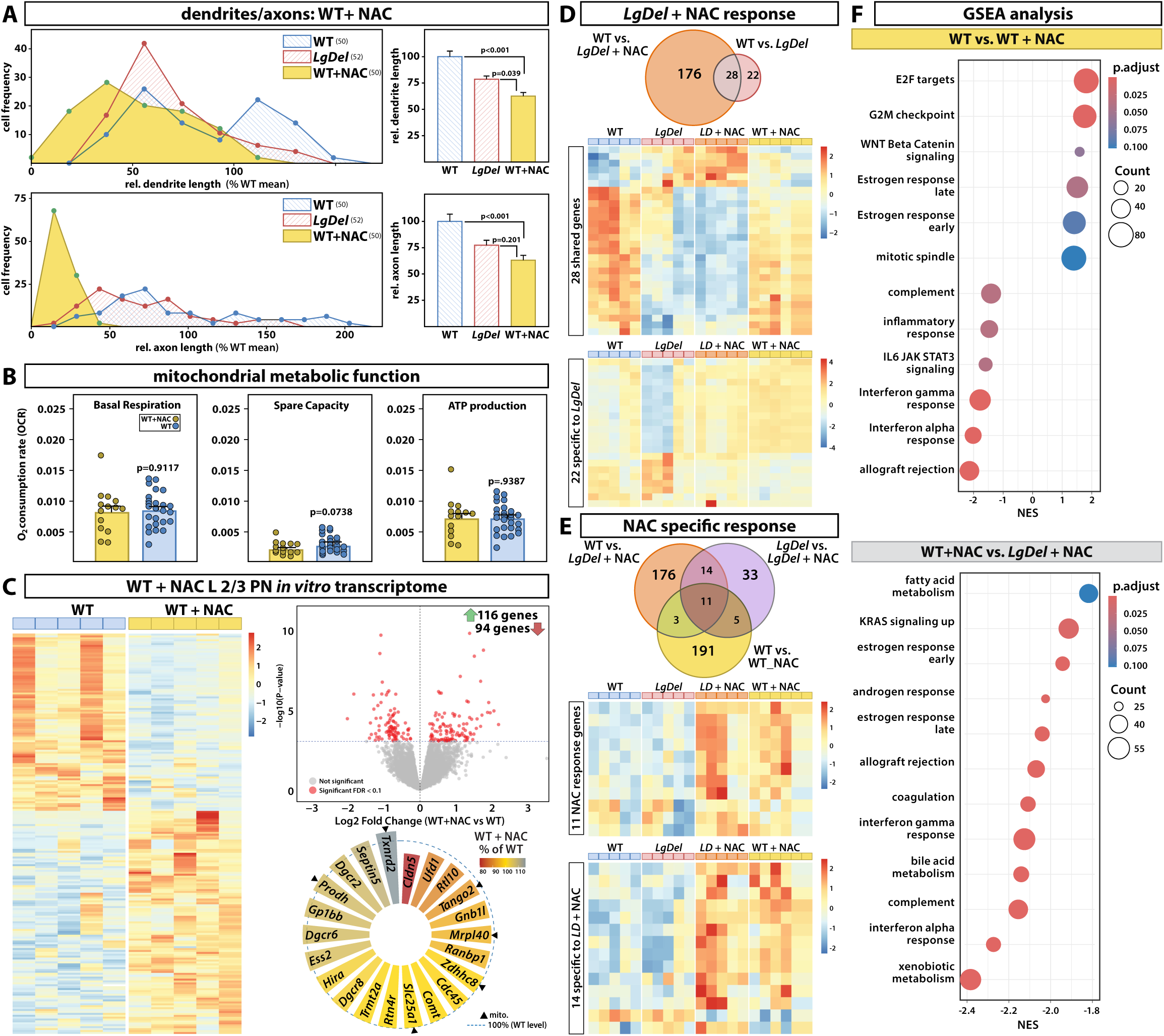
NAC regulated and 22q11 deletion-dependent NAC sensitive genes define a NAC-dependent therapeutic transcriptional response in 22q11-deleted L 2/3 PNs iv. **A.** Distribution of dendrite (top) and axon (bottom) frequencies across WT + NAC L 2/3 PNs *iv*. In both cases, growth is more substantially limited, and the bimodal distribution (dendrites) or “tail” of long axons is no longer apparent (*data re-analyzed from* Fernandez et al, 2019). The mean lengths of WT + NAC dendrites and axons are lower than WT and *LgDel* counterparts (*right*; p = 0.0014 and 0.035; T-test). **B**. Mitochondrial metabolic function (O2 consumption; Agilent Seahorse) in substantially growth-restricted WT L 2/3 PNs treated with NAC remains at levels comparable to both WT as well as *LgDel* (see **Figure 1C**). **C**. Heat maps showing differential expression due to substantial up as well as down-regulation of multiple genes in WT + NAC L 2/3 PNs iv vs. WT counterparts (*left*) (n= 15 fetuses/6 litters WT; 15 fetuses/7 litter WT + NAC). 210 genes are significantly differentially expressed in WT + NAC L 2/3 PNs iv, 116 up- and 94 down-regulated (*top*, *right*). The 22q11-deleted genes, however, are not among this list of DE genes (*bottom*, *right*). **D**. 50 WT vs. *LgDel* L 2/3 PN *iv* DE genes compared to 176 WT vs. *LgDel* + NAC L 2/3 PN *iv* DE genes. Expression of 28 of these DE genes is sensitive to 22q11 deletion and NAC treatment, while another 22 are regulated only by 22q11 deletion, and thus insensitive to NAC. **E**. Comparisons of DE genes identified by integrating WT vs. *LgDel* + NAC, *LgDel* vs. *LgDel* + NAC, and WT vs. WT+ NAC L 2/3 PN iv transcriptomes (top) identifies 11 broadly NAC responsive DE genes (shared across all comparisons) vs. 14 DE genes that respond to NAC only in the context of 22q11 gene deletion in L 2/3 PNs *iv*. Heatmap comparisons confirm NAC-selective transcriptional changes broadly (11 DE genes, *middle*) or selectively for *LgDel* + NAC (14 DE genes, *bottom*). WT + NAC genes, although expressed at similar levels to *LgDel* + NAC are not significantly differentially expressed in the other comparisons. **F**. Hallmark GSEA analysis of WT vs. WT + NAC (top) and WT + NAC vs. *LgDel* + NAC full transcriptomes.

NAC may elicit a distinct transcriptional response that inhibits dendrite and axon growth in WT neurons where ROS declines below typical levels (Mora-Zenil and Moran, 2024; Wilson et al., 2018) vs. that which supports growth in *LgDel* L 2/3 PNs + NAC where elevated ROS returns to WT levels. Thus, we compared the transcriptome of WT + NAC L 2/3 PNs *iv* with WT, *LgDel* and *LgDel* + NAC counterparts (**Figure 4C - F**). Comparison of WT + NAC vs. WT L 2/3 PNs *iv* (**Figure 4C**, *left*) identified 210 DE genes: 116 up- and 94 down-regulated (**Figure 4C**, *top right*). 22q11-deleted genes—including the 6 mitochondrial genes—are not among these DE genes (**Figure 4C**, *bottom right*). To determine whether the 50 DE genes that distinguish *LgDel* L2/3 PNs *iv* from WT are responsive or refractory to NAC, we compared DE genes from WT L 2/3 PNs *iv* with those from *LgDel* + NAC and *LgDel* alone (**Figure 4D**, *top*). Shared differential expression of 28 transcripts common to WT vs. *LgDel* + NAC and WT vs. *LgDel* include 21 22q11-deleted genes and 22 DE genes seen in *LgDel* vs. WT (**Figure 4D**, *middle*). These genes appear to be sensitive to diminished 22q11 gene dosage but not NAC in *LgDel* L 2/3 PNs *iv*.

We compared DE genes across WT vs. *LgDel* + NAC, *LgDel* vs. *LgDel* + NAC, and WT vs. WT+ NAC to distinguish DE genes generally responsive to NAC vs. those that respond to NAC only in the *LgDel* context. 11 DE genes are shared across all three NAC-related pairwise analyses and thus are generally NAC responsive (**Figure 4E**, *top*). These included: the antioxidant defense genes *Srxn1* and *Osgin1*; *Grm2*, a glutamate receptor associated with neuronal activity and growth; *Wif1*, a Wnt inhibitor associated with progenitor proliferation and axon growth, and *Rgs2*, associated with synaptic signaling and plasticity (Brennan et al., 2017; Heo et al., 2006; Hunter et al., 2004; Ingi et al., 1998; Ma et al., 2023; Soriano et al., 2008). These NAC-responsive DE genes were significantly up-regulated in WT and *LgDel* NAC-treated L 2/3 PNs *iv*, while no significant changes were observed in either genotype without NAC (**Figure 4E**, *middle*). In addition, 14 genes are responsive to NAC only in the *LgDel* context (**Figure 4E**, *bottom*). These include 2 genes involved in nucleosome assembly and chromatin remodeling: *Nap1l5* and *Hdac9*, which play essential roles in dendrite growth and synaptic maturation (Attia et al., 2013; Sugo et al., 2010) as well as *Adcyap1*, which encodes the neuropeptide PACAP which can promote neurite outgrowth, and survival (Baskozos et al., 2020). Thus, when DE gene sets are integrated, a distinct set of therapeutic response genes emerges, some generally NAC-responsive, others that respond to NAC only in the context of 22q11 deletion, to support dendrite and axon growth in 22q11-deleted L 2/3 PNs.

GSEA analyses of WT + NAC L 2/3 PNs *iv* (20309 transcripts) vs. WT or *LgDel* + NAC L 2/3 PNs *iv* transcriptomes reinforce the impression that NAC treatment establishes a distinct transcriptional state to support WT L 2/3 PNs *iv* survival in the context of pathologically diminished ROS levels. Enriched gene sets include E2F targets, G2/M checkpoint, and mitotic spindle, all associated with cell survival and growth (**Figure 4F***, top*). Unlike WT vs. *LgDel* or *LgDel* + NAC (see **Figures 1** & **3**), there is no significant enrichment of oxidative metabolism, stress, or mitochondrial sets. There is, however, negative enrichment of gene sets associated with inflammation (**Figure 4F**, *bottom*), consistent with NAC’s anti-inflammatory properties (Askari et al., 2020; Santus et al., 2024). Apparently, *LgDel* L 2/3 PNs iv acquire a unique transcriptional state to support distinct “therapeutic” growth and metabolic state changes.

### Cellular and transcriptional states of 22q11-deleted L 2/3 PNs *in vivo*

NAC reverses L 2/3 PN circuit and related behavioral pathology *in vivo* parallel to its *in vitro* activity (Fernandez et al., 2019). Thus, 22q11-deleted L 2/3 PNs *in vivo* may also acquire a distinct NAC-dependent transcriptional state that supports growth and differentiation similar to that of L 2/3 PNs *in vitro*. To assess L 2/3 PN transcriptional responses that underlie NAC’s therapeutic effects *in vivo*, we established that RNA quantification *in situ* (10X Genomics Xenium) securely identifies L 2/3 PNs and their neighbors so that transcriptional states can be analyzed in intact cortices of early post-natal WT, *LgDel*, *LgDel* + NAC and WT + NAC L 2/3 mice. This approach, though limited in transcript numbers analyzed (347), nevertheless allows reliable identification of all major forebrain cell classes (**Figure 5A**; n = 5 WT, 5 *LgDel*, 5 *LgDel* + NAC, 3 WT + NAC P6 mice; 1 section/mouse analyzed). The relationship between expression levels of key markers, cell type(s), and histological locations is robust across forebrain regions and cell classes, including upper (L 2/3 – 4) vs. lower layer (L 5/6) cortical PNs (**Figure 5B**, **C**). All cell types identified in the pooled UMAP analysis (see **Figure 5A**) are represented in each genotype/NAC treatment (**Figure 5C**, *bottom*); no major cell class is lost or gained in any condition (**Figure 5D**) and proportions are equivalent in each (**Figure 5E**). Finally, a subset of 22q11-deleted genes added to the probe panel appear equally diminished across cell types in *LgDel* or *LgDel* + NAC vs. WT or WT + NAC (**Figure 5F**). Thus, multiplex hybridization/spatial transcriptomic analysis reliably identifies cortical neuron classes for analysis of L 2/3 PN transcriptional changes in response to 22q11 deletion as well as NAC therapy *in vivo*.

**Figure 5:**
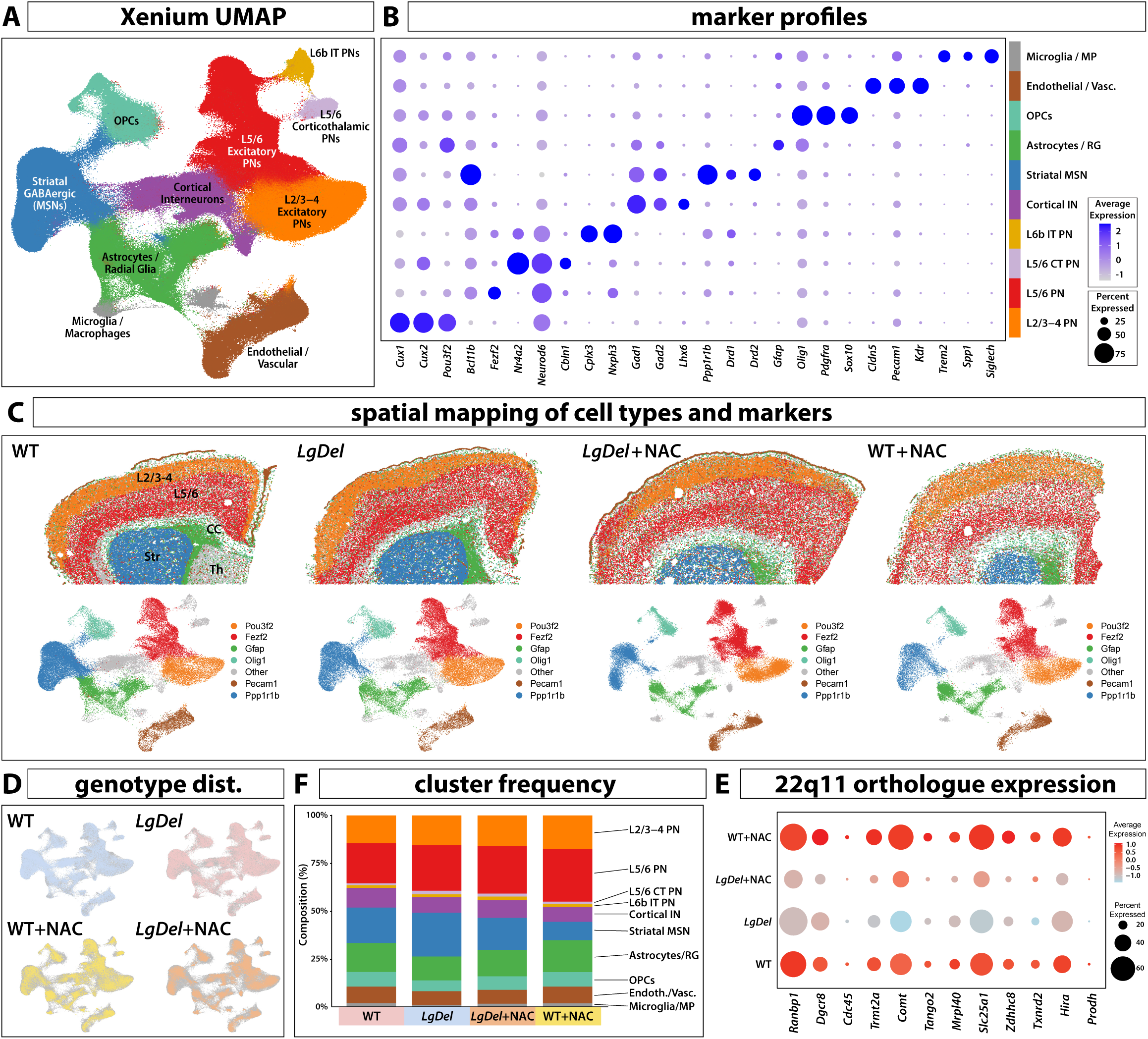
Multiplex quantitative in situ spatial transcriptomic comparison of the WT, LgDel, LgDel + NAC and WT + NAC anterior forebrain, including L 2/3 PNs and their neighbors. **A.** UMAP of forebrain cell types identified by PCA analysis of multiple identity marker genes included in the 350-transcript probe panel. This analysis identifies 10 distinct clusters that reflect identities of known forebrain cell types including L 2/3 PNs and their closely related L 4 counterparts, L 5/6 PNs including two transcriptionally and connectionally distinct sub-classes. **B.** Distribution and magnitude of expression of key marker genes across cell types identified by the unbiased PCA analysis (panel **A**). **C**. Cell types mapped onto representative sections (*top row*) of WT (n = 5 P6 mice, 1 section/mouse), *LgDel* (n = 5 mice, 1 section/mouse), *LgDel* + NAC (n = 5 mice, 1 section/mouse), and WT + NAC (n = 3 mice, 1 section/mouse). The localization of cells classified by PCA is in register with known locations of each cell type. UMAPS for each genotype/treatment condition confirm that no cell classes are lost or gained across the four conditions (*lower row*). **D**. The distribution of all cell clusters in each genotype/treatment condition (colors) correspond to the distribution of all cells analyzed together (gray). **E**. Frequency of cell in each PCA-identified cluster is statistically indistinguishable across the four genotypes/conditions. Proportions of L 2/3-4 vs. L 5/6 cells in *LgDel* and *LgDel* + NAC samples decline modestly, consistent with diminished frequency of L 2/3 PNs due to 22q11 gene deletion (Meechan et al., 2015b; Meechan et al., 2009). **F**. Expression levels of 12 22q11deleted genes included in the 350-transcript probe panel across each of the four genotypes/conditions. Expression varies fairly consistently for these 12 genes across conditions, and their levels are reduced by approximately 50% uniformly in *LgDel* and *LgDel* + NAC.

To determine if growth of individual *LgDel* L 2/3 PN dendrites *in vivo* is limited as *in vitro*, we compared total dendrite length distributions for WT and *LgDel* L 2/3 PNs *in vivo* based upon previously generated data (Fernandez et al, 2019). In contrast with L 2/3 PNs *in vitro*, WT L 2/3 PN *in vivo* dendrite lengths are distributed around a single mean while *LgDel* counterparts are distributed bimodally: a less frequent subset approximates the WT mean, and a more frequent subset falls around a mean 50% of WT (**Figure 6A**, *left*). NAC restores *LgDel* L 2/3 PNs *in vivo* to a broad, flat, unimodal distribution (**Figure 6A**, *middle*), yielding a mean value statistically indistinguishable from WT (**Figure 6A**, *far right*). Unlike growth limiting effects on WT L 2/3 PNs *iv*, NAC has no apparent influence on WT dendrite lengths *in vivo*. Thus, *in vivo* as *in vitro*, diminished 22q11 gene expression changes the distribution of L 2/3 PN dendrite sizes, and NAC enhances growth without returning L 2/3 PN dendrites fully restoring typical distribution.

**Figure 6:**
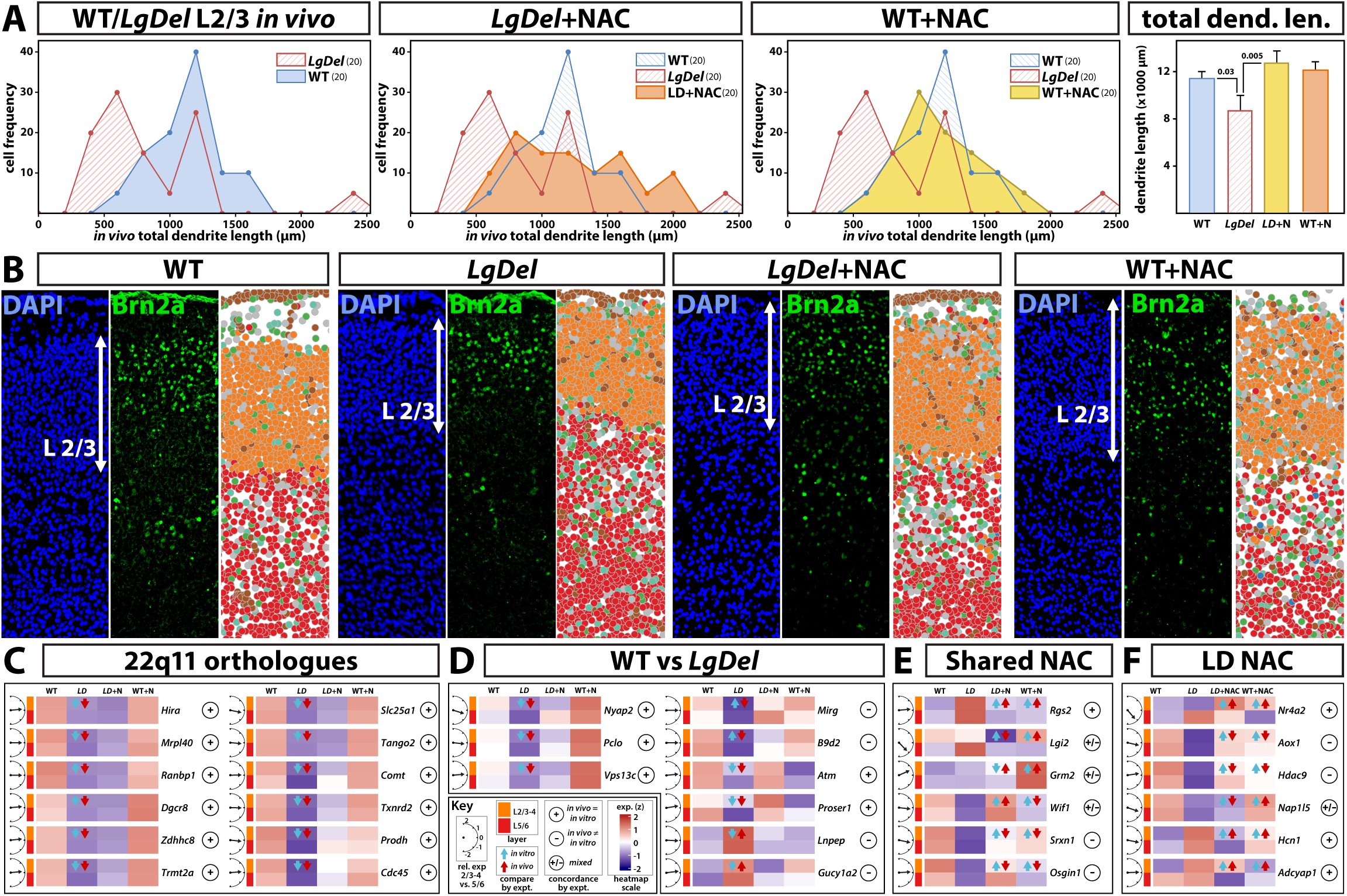
Layer 2/3 PN in vivo transcriptional responses to 22q11 gene deletion and NAC treatment overlap those of Layer 2/3 PNs in vitro. **A.** Distribution of total dendrite length/neuron, measured for WT, *LgDel*, *LgDel* + NAC and WT + NAC L 2/3 PNs labeled *in vivo* by sparse *Cux2*^CreERT^ recombination (*data re-analyzed from* Fernandez et al, 2019). NAC restores *LgDel* mean dendrite length to WT levels, and does not disrupt WT dendrite growth in L 2/3 PNs in vivo (*far right*). **B**. Post-hybridization nuclear and immunohistochemical staining of WT, *LgDel*, *LgDel* + NAC and WT + NAC sections (*left* & *middle* panels) used for spatial transcriptomic analysis (*right* panel) demonstrating integrity of cortical lamination and preservation of selective L 2/3 PN expression of Brn2a, an established marker for subsets of L 2/3 PNs. Diminished thickness and apparent frequency of L 2/3 PNs is seen in both the *LgDel* and *LgDel* + NAC sections, and the corresponding spatial transcriptomic cellular maps. **B.** Uniform diminished expression of a subset of 12 22q11-deleted genes in L 2/3 as well as L 5/6 PNs *in vivo* accords with that *in vitro*. *<u>Inset</u>*: a key for the heatmaps in panels **B** through **E**. **C**. A subset of DE genes from the WT vs. *LgDel* L 2/3 PN *in vitro* comparison are also expressed in L 2/3 PNs *in vivo*, albeit not selectively. **D**. A subset of DE genes identified as generally NACregulated in L 2/3 PNs *in vitro* (i.e. differentially expressed in both LgDel + NAC and WT + NAC L 2/3 PNs *in vitro*) are expressed in vivo. There is enhanced or diminished expression of some of these genes in L 2/3 vs. L 5/6 PNs in vivo and their expression changes in L 2/3 PNs *in vivo* do not fully accord with those *in vitro*. **E**. A subset of DE genes identified as NAC-responsive only in the context of 22q11 deletion. Expression of one gene, *Nr4a2* is substantially enhanced in L 5/6 vs. L 2/3 PNs *in vivo*. L 2/3 PN expression changes *in vivo* for these genes accord generally with those *in vitro*.

To assure that our transcriptome analysis identifies L 2/3 PNs *in vivo* we confirmed identity and location of a molecularly distinct subset by post-hybridization immunostaining (**Figure 6B**). There is an apparent decrease in *LgDel* L 2/3 thickness, and L 2/3 PN frequency consistent with previous observations (Meechan et al., 2009; Meechan et al., 2012; Rukh et al., 2025). Consistent with our *in vitro* data, NAC does not substantially modify diminished L 2/3 or L 5/6 PN expression of 12 22q11-deleted genes the probe set (**Figure 6B**). In addition, the magnitude and direction of L 2/3 PN *in vivo* vs. *in vitro* expression of 4/9 is discordant for 9 *LgDel* vs. WT L 2/3 PN *iv* DE genes (**Figure 6C**). In parallel, NAC also modulates generally NAC-responsive (**Figure 6D**) and *LgDel*-specific NAC-responsive genes (**Figure 6D, E**) in L2/3 PNs *in vivo*; however, the magnitude and direction of *in vitro* vs. *in vivo* expression changes vary. Apparently, expression of 22q11 and DE genes that characterize L 2/3 PNs *in vitro* is preserved *in vivo*; however, quantitative details of expression and response NAC therapy *in vivo* diverge.

### A distinct mitochondria/metabolic transcriptional state in 22q11-deleted L 2/3 PNs in vivo

This *in vitro* and *in vivo* comparison, as well as previous observations (Fernandez et al., 2019) suggest that transcriptional changes in 22q11-deleted L 2/3 PNs primarily mitochondrial and metabolic function. To further evaluate this possibility, we included probes for 25 regulators of mitochondrial integrity, metabolic homeostasis, and antioxidant defense in our spatial transcriptomic analysis. To assess potential differential regulation of these genes, we performed pseudo-bulk analysis in PCA defined L2/3-4 vs. 5/6 PNs (see **Figure 5,6**) to identify WT vs. *LgDel* as well as *LgDel* vs. *LgDel* + NAC DE genes (FDR < 0.05; **Figure 7A, B**). There are considerably more L 2/3-4 PN (73 up- and 123 down-regulated) than L 5/6 PN (28 up- and 70 downregulated) DE genes in the WT vs. *LgDel* comparison. More mitochondrial, metabolic regulatory and antioxidant defense genes are significantly differentially expressed in L 2/3-4 (19/25; **Figure 7A**, *left*) vs. 5/6 (9/25; **Figure 7A**, *right*) PNs *in vivo*, and 15/19 are up-regulated. A similar, but inverted, DE subset emerges for *LgDel* vs. *LgDel* + NAC L 2/3 and 5/6 PNs. 14/25 mitochondrial, metabolic regulatory and antioxidant defense genes are significantly differentially expressed in *LgDel* L 2/3 PNs *in vivo* in response to NAC (**Figure 7B**, *left*), and 9/25 are differentially expressed in L 5/6 PN counterparts (**Figure 7B**, *right*). Finally, some variation in expression of mitochondria, metabolic regulatory and antioxidant defense genes—none of which are DE genes *in vitro*—is seen in L 2/3 PNs *in vitro*; however, direction and magnitude of expression differences diverge (**Figure 7C**). Apparently, NAC selectively reverses L 2/3 PN mitochondrial metabolic regulatory, and antioxidant defense gene expression changes due to 22q11 deletion *in vivo* as *in vitro* without altering expression levels of 22q11-deleted or downstream DE genes; however, *in vivo* responses differ quantitatively from those in L 2/3 PNs *in vitro*.

**Figure 7:**
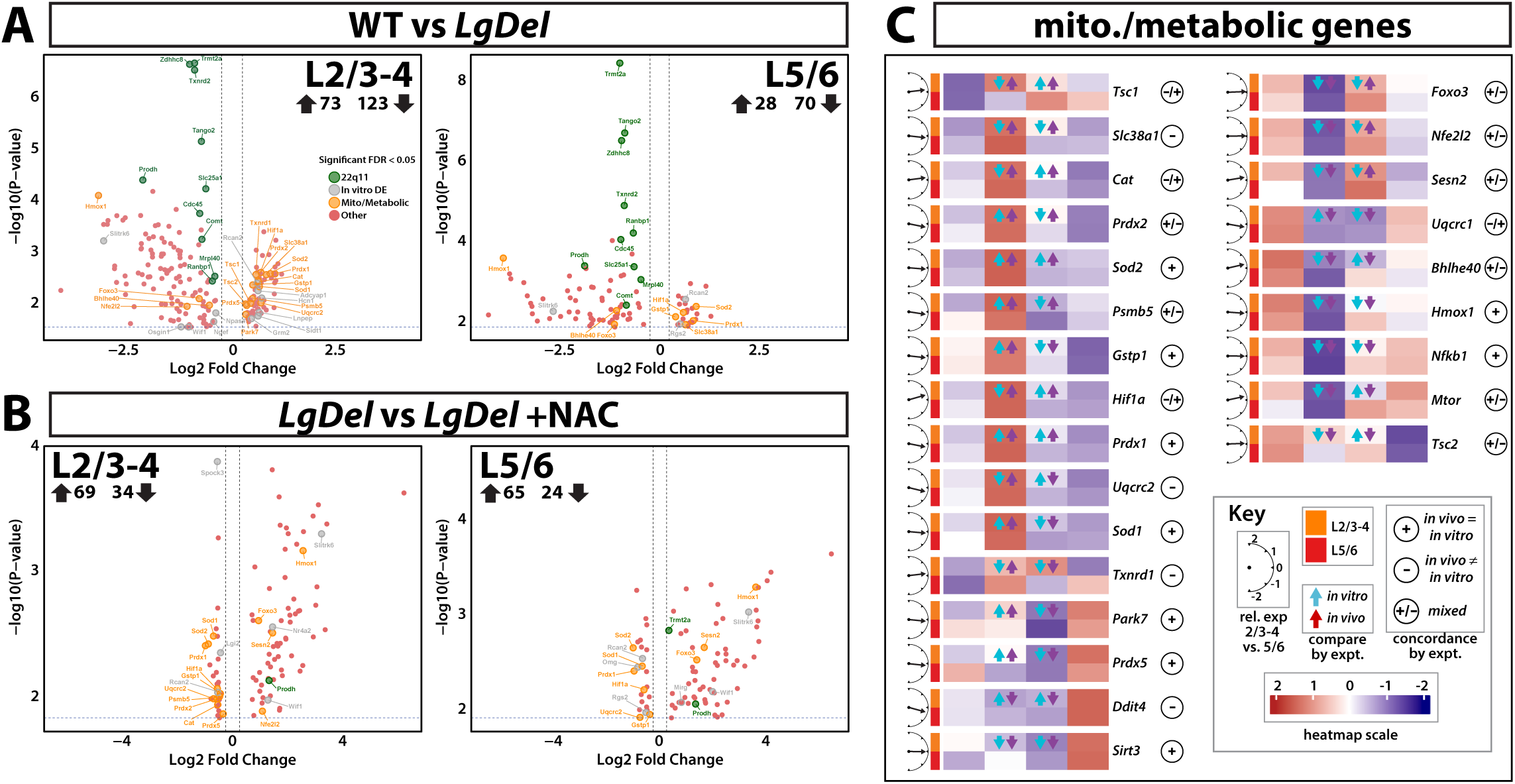
22q11-deleted L 2/3 PNs in vivo have a mitochondria/metabolic transcriptional state distinct from L 2/3 PN in vitro counterparts or L 5/6 PN in vivo neighbors. **A**. Volcano plots of significantly differentially expressed (DE) transcripts based upon comparison of transcript levels via pseudo-bulk seq analysis for individual L 2/3 (*left*) vs L 5/6 PNs *in vivo* (*right*) from spatial transcriptomic data described above. In each plot down- (*left*) and upregulated DE genes (*right*) from four key categories are shown: 22q11-deleted genes (green dots), L 2/3 PN *in vitro* DE genes (gray dots), mitochondrial/metabolic regulatory genes, including antioxidant defense, mitochondrial biogenesis and maintenance genes (orange dots) and additional genes in the probe panel (pink dots). **B**. Volcano plots of significant DE genes in a pseudo-bulk seq comparison of *LgDel* vs. *LgDel* + NAC L 2/3 (*left*) vs. L 5/6 PNs (*right*) *in vivo*. The four categories of DE genes are indicated as in panel **A**. **C**. Comparison of L 2/3 PN *in vivo* vs. *in vitro* as well as L 5/6 PNs *in vivo* expression levels of the full set of 25 mitochondrial and metabolic regulatory genes chosen to assess oxidative stress, antioxidant defense, and their modulation by NAC. Relative expression levels in L 2/3 vs. 5/6 PNs *in vivo*, as well as concordance or divergence of expression for each gene is indicated as in Figure 6.

## DISCUSSION

NAC antioxidant therapy reverses oxidative stress-associated developmental pathology due to diminished dosage of 22q11-deleted genes in a specific neuron class—cerebral cortical L 2/3 PNs by circumventing apparently irreversible changes in modes of dendrite and axon growth as well as differential gene expression. NAC does not substantially enhance expression of 22q11 orthologues nor modify 22q11 deletion-dependent downstream transcriptional changes in L 2/3 PNs *in vitro* or *in vivo*. Instead, NAC engages novel antioxidant defense mechanisms that support divergent, but beneficial, dendrite, axon, and synapse growth and differentiation via 22q11 gene dosage-independent pathways. General features of this therapeutic response are recognized in primary cultured 22q11-deleted L 2/3 PNs; however, *in vitro* vs. *in vivo* cellular and transcriptional changes diverge substantially. Thus, therapeutic mechanisms assessed *in vitro*— even in primary cultured neurons that retain several aspects of *in vivo* identity—likely require *in vivo* validation to define precise molecular pathways underlying therapeutic responses.

The loss of bimodally distributed dendrite and axon sizes in 22q11-deleted L 2/3 PNs *iv*; the inversion of this relationship *in vivo*, and failure of NAC to fully restore WT distributions *in vitro* or *in vivo* raises three related questions: First, does diminished 22q11 gene dosage foreclose a “maximal” L 2/3 PN growth program? Second, does this foreclosure reflect irreversible consequences of oxidative stress in differentiating L 2/3 PNs? Third, could loss of maximal growth reflect altered genesis of L 2/3 PN neuroblasts (Meechan et al, 2009, 2015; Rukh et al, 2025) that eliminates a sub-population capable of maximal dendrite and axon growth? If 22q11 deletion-dependent oxidative stress disrupts L 2/3 PN progenitor proliferation (Campbell et al., 2023; Paranjape et al., 2023; Purcell et al., 2023), diminishing ROS or enhancing antioxidant defense during L 2/3 PN neurogenesis could potentially restore maximal growth by stabilizing genesis of neuroblasts capable of maximal growth. Alternately, if L 2/3 PN neuroblasts capable of maximal growth are lost due to divergent neurogenesis independent of oxidative stress, antioxidant therapy alone, via NAC or other antioxidants, would likely not restore the WT distribution of L 2/3 PN dendrite or axon arbors.

We showed previously that <u>acutely</u> diminished *Txnrd2* expression *in vitro* or conditional heterozygous deletion *in vivo* in differentiating L 2/3 PNs recapitulates key aspects of L 2/3 PN pathology due to broader 22q11 deletion (Fernandez et al, 2019). In contrast, <u>constitutive</u> *Txnrd2* heterozygous deletion does not yield cellular, metabolic, or transcriptional changes in L 2/3 PNs *iv*, making it unlikely that it acts as a “contiguous gene” to drive in isolation key L 2/3 PN growth phenotypes associated with broader 22q11 gene deletion. The WT bimodal distribution of dendrite and axon arbor sizes remains despite diminished growth of larger and smaller *Txnrd2*^+/-^ L 2/3 PN subpopulations. Moreover, *Txnrd2* heterozygous mutation, unlike broader 22q11 deletion diminishes mitochondrial metabolic activity. Apparently, constitutively diminished *Txnrd2* dosage has a distinct and potentially cumulative influence on L 2/3 PN growth and mitochondrial/metabolic function *in vitro* that is neither like that in WT nor broader 22q11 gene deletion. Indeed, physiological rather than transcriptional mechanisms may preserve L 2/3 PN viability *in vitro* by uniformly limiting growth, perhaps via diminished mitochondrial oxidative phosphorylation. The extent to which antioxidants reverse L 2/3 PN metabolic pathology associated with constitutive heterozygous *Txnd2* mutation and its relevance to parallel responses in 22q11-deleted L 2/3 PNs remains to be determined.

Based upon transcriptional changes, NAC *in vivo* modulates L 2/3 PN homeostatic mechanisms not directly compromised by 22q11 gene deletion—a distinct set of genes is NAC responsive while those sensitive to 22q11 dosage are unchanged. Apparently, the ROS-scavenging activity of NAC cannot reverse direct transcriptional consequences of diminished dosage of multiple 22q11 orthologues expressed in differentiating L 2/3 PNs *in vitro* or *in vivo*. Nevertheless, NAC, presumably by diminishing ROS and reducing oxidative stress in L 2/3 PNs (Fernandez et al, 2019), elicits a novel transcriptional response that circumvents rather 22q11deletion dependent gene expression changes to support therapeutically beneficial— but divergent—L 2/3 PN growth and differentiation (**Figure 8**). Thus, identifying physiological states *in vitro* or *in vivo* underlying specific NDD developmental pathology may provide opportunities to exploit flexible cellular mechanisms and transcriptional networks as an alternative to direct restoration of function of mutant genes or their downstream targets.

**Figure 8:**
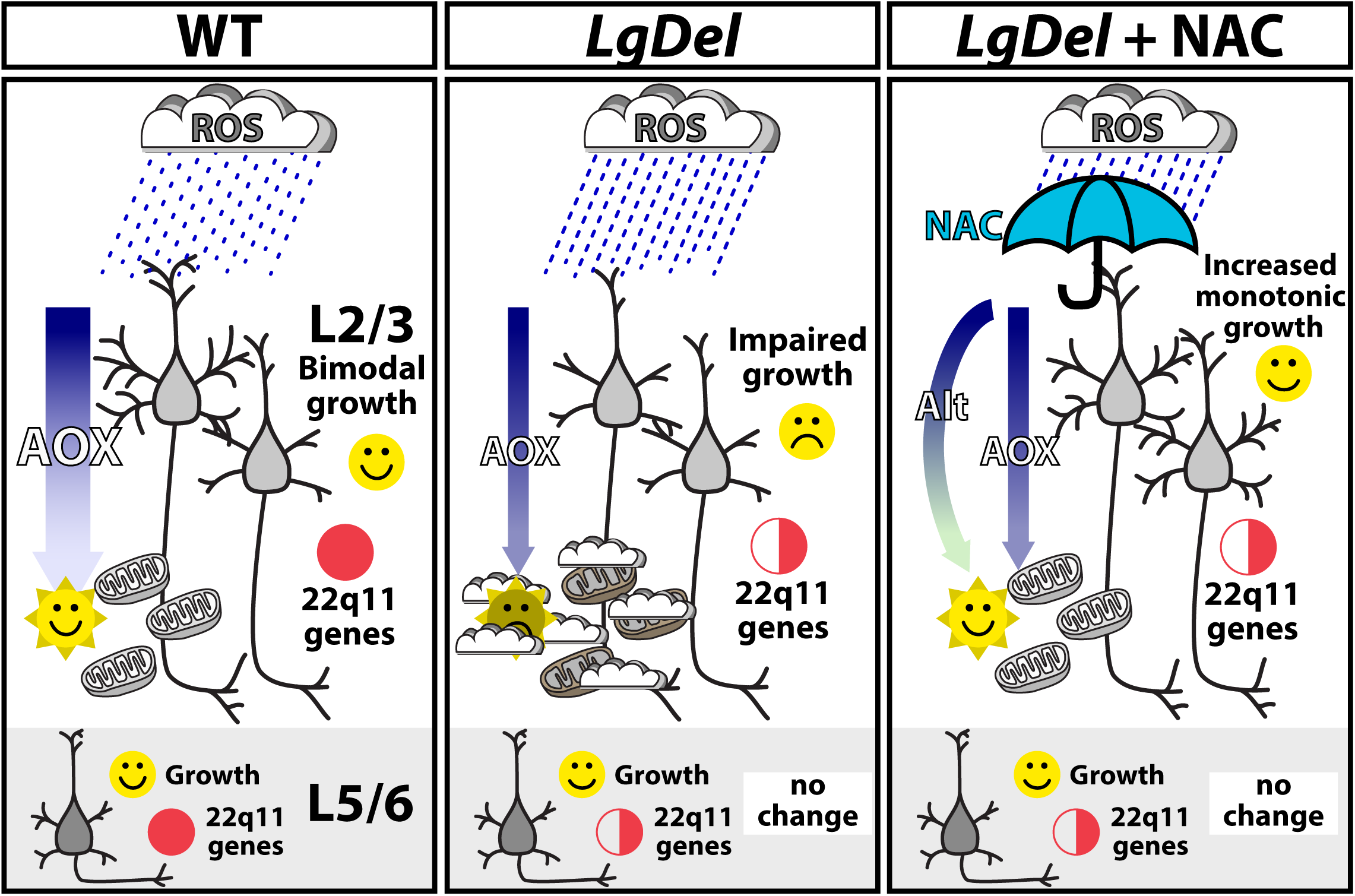
NAC therapy selectively circumvents oxidative stress-mediated restricted growth in 22q11-deleted L 2/3 PNs by engaging distinct cellular and transcriptional responses. ***<u>Left</u>:*** L 2/3 PNs in the developing cortex have a range of sizes, and a unique metabolic/ antioxidant defense transcriptional signature that makes them differentially sensitive to oxidative stress. L 5/6 PNs do not share this distinct metabolic/antioxidant defense signature. *<u>Middle</u>*: 22q11 gene deletion limits L 2/3 PN growth, especially that which approximates the largest WT L 2/3 PNs, and leads to mitochondrial changes due to increased ROS and resulting oxidative stress. These changes are not seen in L 5/6 PNs, despite diminished expression of 22q11 genes in these neurons. ***<u>Right</u>:*** NAC anti-oxidant therapy, *in vitro* or *in vivo*, modifies the diminished growth of L 2/3 PNs without returning them to a WT distribution of dendritic arbor sizes. This distinct therapeutically beneficial cellular response is limited to L 2/3 PNs *in vivo*, and is accompanied by a transcriptional response that neither restores expression of 22q11 genes from their 50% decrement, nor any of the downstream genes that appear to change their expression in response to diminished 22q11 gene dosage. Instead, an alternate transcriptional response includes novel antioxidant defense genes that presumably support a divergent but salutary increase in L 2/3 PN growth and differentiation.

Our *in vivo* analysis suggests that 22q11-deleted L 2/3 PNs are in a unique local oxidative environment during early post-natal association cortico-cortical circuit development *in vivo*. This environment and related NAC-mediated transcriptional changes is not replicated in 22q11deleted L 5/6 PNs *in vivo*, despite shared diminished expression of multiple 22q11 deleted genes. These results reinforce the conclusion that laminar-selective oxidative stress in L 2/3 PNs and perhaps some of their interneuron and glial neighbors (Lee et al., 2021; Shin et al., 2024; Steullet et al., 2017; Wang and Michaelis, 2010) is a significant driver of association corticocortical circuit pathology as well as its functional and behavioral consequences in the *LgDel* 22q11DS mouse model. This L 2/3 selective pathology may emerge only as demands for bioenergetic support of growth and synaptogenesis increase during early post-natal life (Attwell and Laughlin, 2001; Chugani, 1998; Fernandez et al., 2019), or may have antecedents during initial L 2/3 PN genesis. Divergent proliferative and neurogenic capacities of 22q11-deleted L 2/3 PN intermediate progenitors alter frequency and identities of L 2/3 PNs (Fenelon et al., 2013; Marinaro et al., 2017; Meechan et al., 2009; Paronett et al., 2015; Rukh et al., 2025). In contrast, L 5/6 PNs and the apical progenitors from which they are primarily derived seem more resilient to 22q11 deletion, despite diminished expression of the same 22q11-deleted genes. It remains to be determined if selective L 2/3 vulnerability to metabolic stress in multiple NDDs is a shared pathogenic mechanism broadly sensitive to mitochondria-targeted therapies interventions (Frye et al., 2024; Khan et al., 2020; Pinto Payares et al., 2024; Rojas-Charry et al., 2021)

Our results indicate that *in vitro* assays relying upon patient-derived or engineered induced pluripotent stem cells (IPSCs) with 22q11 deletions (Liang and Zhang, 2013; Yoshihara et al., 2017) may require extensive adjustment to reliably identify pathogenic mechanisms and therapeutic targets. Variability of phenotypes reported in several recent studies using engineered or patient-derived IPSC planar cultures or cortical organoids carrying 22q11 deletions amplify concerns regarding these technical challenges (Rao et al., 2025b; Tegtmeyer et al., 2025). In most IPSC-based assays—planar cultures or organoids—some aspects of cellular and transcriptional identity are preserved; however, challenges including oxidative stress complicate alignment of *in vitro* and *in vivo* neuron classes (Bhaduri et al., 2020; Castiglione et al., 2025; Uzquiano et al., 2022). For 22q11DS, adjacent cortical neurons or glia not available *in vitro*, as well as 22q11 deletion-dependent systemic cardiovascular or immune changes (Cioffi et al., 2022; Crockett et al., 2021) may contribute to a uniquely aberrant L 2/3 metabolic environment *in vivo* as a foundation for beneficial therapeutic responses not suggested by *in vitro* assays. Understanding the balance of cell autonomy *in vitro* and systemic influences *in vivo* is key to optimize therapeutic approaches for a broad range of polygenic NDD pathologies (Eyring and Geschwind, 2021; LaMantia, 2024; Wang et al., 2022). Accordingly, *in vivo* validation of suggestive *in vitro* findings—especially in genomically valid NDD animal models—is imperative to fully exploit pathological mechanistic insights as a reliable foundation for effective NDD therapies.

## MATERIALS and METHODS

### Animals

Mice carrying a hemizygous deletion on chromosome 16 (*Idd* to *Hira*: “*LgDel”*) were maintained on a C57Bl/6N background. To minimize genetic drift, *LgDel* males were routinely crossed with C57Bl/6N wild-type (WT) females obtained from Charles River Laboratories. Timed matings were established by pairing *LgDel* males with WT females overnight; morning of vaginal plug detection was designated as embryonic day (E) 0.5. Pregnant dams were euthanized by rapid cervical dislocation at E16.5, and fetuses were collected in RNase-free HEPES buffer under sterile conditions for downstream processing. For postnatal experiments, natural deliveries were monitored daily, and the day of birth was recorded as postnatal day (P) 0 and P6 pups were collected for analysis. For genotyping, tail or limb biopsies were obtained from each embryo or pup. Samples were digested overnight at 60°C in SNET buffer supplemented with Proteinase K. Genomic DNA was extracted using phenol/chloroform precipitation and used for PCR based genotyping to identify WT and *LgDel* genotypes.

### Electroporation and Primary Neuronal Culture

E16.5 embryos were collected from timed pregnant litters and dorsal telencephalic progenitors/early neuroblasts were transfected with pCI-GFP plasmid DNA injected into the lateral ventricles (Fernandez et al., 2019; Hand et al., 2005). Following electroporation, cortices were dissected, dissociated (Worthington Biochemical, USA), and cultured for five days *in vitro* (DIV) as described (Ahlemeyer and Baumgart-Vogt, 2005; Fernandez et al., 2019). After 5 DIV, neurons exhibiting at least two distinct dendrites and a single axon measuring at least twice the length of the longest dendrite were selected for analysis. Neurons were reconstructed and traced (Imaris) to quantify total dendritic and axonal length as well as branch numbers and branching order.

### NAC treatment in vitro and in vivo

N-acetylcysteine (NAC; PharmaNAC) was administered at a final concentration of 1 mM during cell plating and replenished with each media change at 24 hours (1 DIV) and 96 hours (4 DIV). For in vivo experiments, NAC was delivered to pregnant dams by supplementing the drinking water with 900 mg/L NAC from embryonic day 14. The pups continued to receive NAC in their drinking water after birth at the same concentration (900 mg/L), with fresh solutions prepared three times per week until tissue collection at P6 (Cabungcal et al., 2013; das Neves Duarte et al., 2012; Fernandez et al., 2019).

#### Seahorse metabolic analysis

Primary cortical neurons were dissociated from embryonic brains (E16.5) and plated at ∼140,000 cells per well as described above. Cultures were maintained with standard media changes before being switched to XF DMEM Seahorse media for metabolic assays. After a twostep equilibration, oligomycin, FCCP, and rotenone/antimycin A were loaded per manufacturer’s (Agilent) guidelines, and OCR was recorded using the Wave platform.

### RNA sequencing

RNA was extracted from DIV5 primary cultures of WT and *LgDel*, Txnrd2^+/-^, WT+NAC, *LgDel*+NAC groups. For each condition, isolated RNA with integrity (RIN) scores of 10 (Agilent RNA Tapestation) were pooled to generate minimum five biological replicates each condition. Libraries were generated and sequenced on an Illumina HiSeq platform (Genewiz), yielding at least depth of 90 million paired end 150bp long reads per sample. Reads quality was assessed with FastQC, and trimming/filtering was performed with BBDuk. Reads were aligned to the GRCm38/mm10 genome using STAR, achieving a minimum unique alignment rate of 85%. Transcript abundance was quantified as TPM. Differential gene expression analysis was performed using edgeR in R. Low count genes (<15 reads in ≥3 samples) were filtered out. Data were normalized, and a robust dispersion estimation was used. Differentially expressed genes (DEGs) were identified using a likelihood ratio test with an FDR cutoff of < 0.1. GSEA analysis was performed using fgsea function in R (Korotkevich et al., 2021). Gene ontology enrichment for DE genes was performed in Metascape using default settings (Zhou et al., 2019).

### Histological preparation

P6 pups were collected, rinsed in PBS, and immersion fixed in 4% paraformaldehyde overnight. Following cryoprotection with a gradient of 10%, 20%, and 30% sucrose in PBS, embryos were embedded in OCT in a consistent orientation and stored in −80°C. 12µm coronal sections were cut on a cryostat from OCT embedded embryos and slides were dried at RT and stored in −80°C.

### Real-Time quantitative (q)PCR

cDNA synthesis reactions were conducted as previously described (Maynard et al., 2013) on RNA isolated from primary cultures. qPCR for multiple transcript-specific primers was performed on a CFX384 thermal cycler using EvaGreen; BioRad.

### Xenium Spatial Transcriptomic Analysis

Fresh frozen tissue sections of 12um thickness stored at −80°C were processed according to 10x standard procedure (Janesick et al., 2023). Xenium spatial sequencing data were analyzed using the 10x Genomics Xenium platform and subsequent processing was performed in R (v4.5) with the Seurat (v5.1.0) and SeuratDisk packages (Butler et al., 2018). Raw outputs were imported via LoadXenium() and filtered for low quality cells based on number of transcripts (nCount), number of expressed genes (nFeature). Expression counts were log-normalized using Seurat’s NormalizeData() function, and variable features were identified via FindVariableFeatures (Butler et al., 2018) using the “vst” method and canonical cell type markers to annotate clusters (Zeisel et al., 2015). Dimensionality reduction was performed by Principal Component analysis followed by uniform manifold approximation and projection (UMAP) for visualization. The dataset was clustered using the Louvain algorithm (FindNeighbors() and FindClusters()) at a resolution of 0.1, resulting in 10 transcriptionally distinct clusters. Spatial visualization was generated using the SpatialDimPlot() and SpatialFeaturePlot() functions, overlaying cluster or gene expression information on the section image. Additional spatial analyses, including expression plots and marker gene identification, were performed using FindAllMarkers() with the Wilcoxon rank-sum test. Cell proportional analysis was performed using ANOVA with Dunnett’s post hoc test, comparing all conditions to WT controls. DE analysis of 347 genes across all conditions was performed using EdgeR in R (Robinson and Smyth, 2007). All analyses were performed on a high-performance workstation (Intel-Xeon Silver 4208 CPU, 16 threads, 256 GB RAM).

## ACKNOWLEDGEMENTS

We thank Elizabeth Kunst, Grace Boyer, and Hannah Young for technical assistance. This work was supported by grants to ASL from the National Institute of Mental Health (MH126294), the National Institute of Child Health and Human Development (HD042182) and the Red Gates Foundation.

