## Supplemental Figure 1. for "Distinct cellular and transcriptional mechanisms mediate an antioxidant therapeutic response in 22q11-deleted upper layer cortical projection neurons"

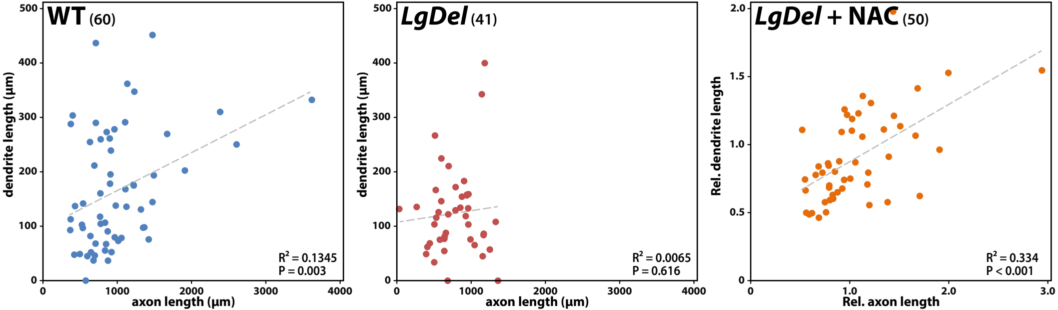


Supplemental Figure 1: Correlations between dendrite and axon sizes per cell for WT, LgDel, LgDel + NAC L 2/3 PNs in vitro. These correlograms are based upon data plotted for dendrite and axon sizes in main Figures 1 and 3.
